# Silhouette width using generalized mean – a flexible method for assessing clustering efficiency

**DOI:** 10.1101/434100

**Authors:** Attila Lengyel, Zoltán Botta-Dukát

## Abstract

1. Cluster analysis plays vital role in pattern recognition in several fields of science. Silhouette width is a widely used measure for assessing the fit of individual objects in the classification, as well as the quality of clusters and the entire classification. This index uses two clustering criteria, compactness (average within-cluster distances) and separation (average between-cluster distances), which implies that spherical cluster shapes are preferred over others – a property that can be seen as a disadvantage in the presence 22 of clusters with high internal heterogeneity, which is common in real situations.
2. We suggest a generalization of the silhouette width using the generalized mean. By changing the *p* parameter of the generalized mean between −∞ and +∞, several specific summary statistics, including the minimum, maximum, the arithmetic, harmonic, and geometric means, can be reproduced. Implementing the generalized mean in the calculation of silhouette width allows for changing the sensitivity of the index to compactness vs. connectedness. With higher sensitivity to connectedness instead of compactness the preference of silhouette width towards spherical clusters is expected to reduce. We test the performance of the generalized silhouette width on artificial data sets and on the Iris data set. We examine how classifications with different numbers of clusters prepared by single linkage, group average, and complete linkage algorithms are evaluated, if *p* is set to different values.
3. When *p* was negative, well separated clusters achieved high silhouette widths despite their elongated or circular shapes. Positive values of *p* increased the importance of compactness, hence the preference towards spherical clusters became even more detectable. With low *p*, single linkage clustering was deemed the most efficient clustering method, while with higher parameter values the performance of group average and complete linkage seemed better.
4. The generalized silhouette width is a promising tool for assessing clustering quality. It allows for adjusting the contribution of compactness and connectedness criteria to the index value, thus avoiding underestimation of clustering efficiency in the presence of clusters with high internal heterogeneity.

## Introduction

Cluster analysis is the method of grouping similar objects in order to simplify the structure of a data set. It is concerned with discontinuous variation in the data set, that allows for separating and identifying ‘types’ of objects. Clustering is a common exploratory tool for pattern recognition in large samples in various fields of science, like geoinformatics (e.g. Lu et al. 2016), genomics (e.g. Ramoni et al. 2002), epidemiology (e.g. Kenyon et al. 2014), or psychology (e.g. Clatworthy et al. 2005). Moreover, classification is a prerequisite for naming abstract entities like biogeographical regions and habitat types, thus it is a basic statistical approach in bioregionalization (e.g. González-Orozco et al. 2013, Lechner et al. 2016), and vegetation typology on different scales (e.g. De Cáceres et al. 2015, Lengyel et al. 2016, Marcenò et al. 2018). Clustering methods are often divided into crisp and fuzzy methods (Podani 2000). Crisp clustering procedures provide unequivocal assignment of objects to groups, while fuzzy methods express memberships as weights. The advantage of fuzzy classification over crisp methods is that they allow for differentiation of typical, transitional, and outlier objects (De Cáceres et al. 2010). However, fuzzy algorithms are much more intensive computationally and they require more subjective decisions from the user for the parameterization; therefore, crisp methods are still the most widespread. Crisp classifications can be further divided into hierarchical and non-hierarchical methods on the condition whether they classify the objects into a groups which are nested subsets of each other or a simple partition without nested structure.

By its basically descriptive nature, clustering techniques, especially crisp algorithms, produce classifications even if there is no discontinuity in the data set, potentially leading to false conclusions about the within-sample variation. A plethora of methods is available for testing the quality (also called validity or efficiency) of classifications, each applying more or less differently formalized criteria (Milligan & Cooper 1985, Handl et al. 2005; Vendramin et al. 2010). One of the most commonly applied methods is silhouette width (Rousseeuw 1987), which encompasses two clustering criteria: *separation* (i.e., average distance between objects of different clusters) and *compactness* (i.e., average within-cluster distance) (Handl et al. 2005). Silhouette width is calculated for each object of the classification thus indicating how well they fit into their respective cluster. The cluster-wise or the global mean of objects can be used to assess the distinctness of specific clusters or the validity of the total classification, respectively. Due to the compactness criterion involved as average within-cluster distance, silhouette prefers spherical cluster shapes (Rousseeuw 1987); however, in practice clusters can possess different shapes according to their structure in the multidimensional space of the variables. Moreover, each clustering algorithm has its own tendency to produce clusters with certain characteristics, including cluster shape, and evaluating them by validity indices following different shape criteria can bring misleading results (Handl et al. 2005). Hence, in the presence of non-spherical clusters, silhouette width may falsely suggest low classification efficiency. Those indices are more suitable for elongated or irregular cluster shapes which quantify the degree to which objects are assigned to the same cluster as their nearest neighbours, i.e. those applying the *connectedness* criterion (Saha & Bandyopadhyay 2012).

In this paper we propose a generalization of the silhouette width. Applying the generalized mean which gives adjustable mean ranging between the minimum and maximum, we propose a flexible formula which allows for scaling the sensitivity of the index between connectedness and compactness, thus allowing high values for non-spherical clusters. This enables users to optimize classifications for different cluster shapes. The use of the new method is illustrated on artificial point patterns and a widely known real sample data set.

## Materials and Methods

### The original silhouette width

The original definition of silhouette width according to Rousseeuw (1987) is as follows. Let *i* be a focal object belonging to cluster *A*. Denote by *C* a cluster not containing *i*. *a*(*i*) is defined as the average dissimilarity between *i* and all other objects in *A*, while *c*(*i*,*C*) is the average dissimilarity between *i* and all objects in *C*.

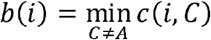

The silhouette width, *s(i)*, is defined as:

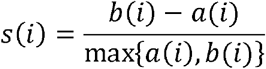

*s(i)* ranges between −1 and 1. Values near 1 indicate that object *i* is much closer to the other objects in the same cluster than to objects of the second closest cluster, implying a correct classification. If *s(i)* is near 0, the correct classification of the focal object is doubtful, thus suggesting intermediate position between two clusters. *s(i)* near −1 indicates obvious misclassification. Accordingly, averaging silhouette widths over a cluster gives an assessment of the ‘goodness’ of that cluster, or a sample-wise average can be used as an index of the validity of the entire classification. Instead of cluster-wise or sample-wise averages of *s(i)*, the number or proportion of objects with positive silhouette width can also be used as validity measures. For a cluster containing a single object, *s(i)* takes the arbitrary value 0.

### Implementing the generalized mean

Applying the arithmetic mean to calculate average within- and between-cluster distances, as the index was introduced originally (Rousseeuw 1987), implies that the ideal cluster shape is spherical. However, this preference can be relaxed by choosing other types of means. Generalized mean (also called Hölder or power mean) offers a flexible solution to calculate sample means ranging between minimum and maximum (Cantrell & Weisstein, http://mathworld.wolfram.com/PowerMean.html). Let *X* be a sample of positive real numbers *x*_*1*_, *x*_*2*_, …, *x*_*n*_ and *p* an element of affinely extended real numbers. The generalized mean of degree *p* is:

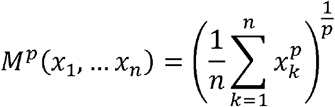

For *p* = 0 and *p* = |∞| the following exceptions are to be made:

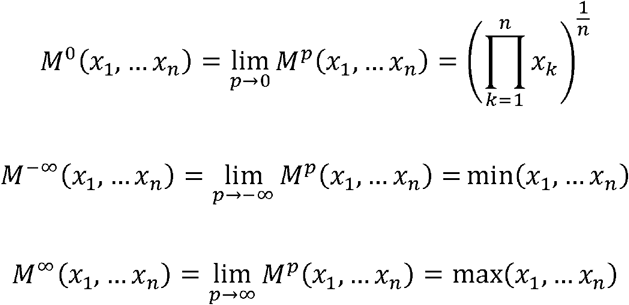

The generalized mean takes the values of well-known summary statistics presented in Table 1. The original version of silhouette width is the special case when within- and between-group average distances are calculated by *p* = 1. By changing the *p* parameter it is possible to emphasize lower or higher distances in the calculation of means. The lower the *p* value is, the more importance is attributed to objects in close proximity, while the effect of farther neighbour objects (including outliers) is reduced. In this way, the criteria of compactness is gradually replaced by connectedness and clusters with irregular or elongated shape can also be considered ‘good’. At *p* = −∞ a classification is ideal if each object is assigned to the same cluster as the most similar other object in the sample. This procedure follows the logic of single linkage clustering, while the original version making use of arithmetic averages followed the logic of average linkage. In contrast, when *p* > 1, the compactness criterion is attributed higher weight, thus the preference towards spherical clusters is further increased and the effect of outliers on the overall classification should become more significant. At *p* = +∞ the clustering criteria of complete linkage is applied.

### Data sets and tests

We test the performance of the generalized mean with different parameterization on artificial point patterns and well-known public data sets.

**Table 1.** Special cases of the generalized mean

| $p$ | *descriptive statistic* |
| --- | --- |
| $-\infty$ | minimum |
| -1 | harmonic mean |
| 0 | geometric mean |
| 1 | arithmetic mean |
| 2 | quadratic mean (root-mean-square) |
| $+\infty$ | maximum |

Artificial data sets containing 100 objects and two variables were generated. The data sets represented data structures some of which were also applied by Podani (2000) for the illustration of the behaviour of different clustering methods: 1) completely random point pattern without true clustered structure, points on the two sides of the plane are assigned to different clusters (both separation and compactness are low); 2) two clusters with few transitional elements between them (moderate separation and compactness); 3) four distinct point aggregations corresponding to four clusters (high separation, high compactness); 4) the same four clusters but only two clusters are defined, each comprising two aggregations (high true separation and compactness but too low number of clusters); 5) two well separated clusters of unequal size (20 vs. 80 points) and spread (high separation, high compactness, unequal size); 6) three clusters of elongated shape running parallel, well separated, but heterogeneous clusters (high separation, low compactness); 7) two concentric clusters (high separation, different compactness, special spatial arrangement). The analyses were run also with randomly permuted group memberships on all the above point patterns in order to test the performance of generalized silhouette width in the case of inefficient clustering but here we show only the results with data set 3 (i.e., four distinct point aggregations and four clusters).

The Iris data set was originally published by Fisher (1936). It contains morphological measurements of 150 individuals of *Iris setosa*, *I. virginica*, and *I. versicolor*, 50 individuals each. *I. setosa* is morphometrically distinctly separated from the other two, while *I. virginica* and *I. versicolor* differ rather gradually. The original data set contained four variables, from which we used only two, sepal length and petal length. Species assignment was used as a priori classification. Data was accessed from the vegan (Oksanen et al. 2018) package of the R software (R Core Team 2017), then variables were standardized to mean = 0 and standard deviation = 1.

On these data sets generalized silhouette widths with different *p* parameter values were calculated using the a priori classifications. The patterns of misclassified objects on the point scatters were assessed visually. Overall classification quality was measured by misclassification rate (MR; the number of misclassified objects in the sample divided by the total number of objects) and mean silhouette width (MSW; the sample-wise mean of *s(i)*).

We evaluated also the performance of different classification methods in the view of the generalized silhouette width. For this purpose, we used a two-dimensional random point pattern of 1000 points because we supposed that in the lack of true cluster structure the inherent characteristics of the methods will determine classification the most. We classified this data set using single linkage, group average and complete linkage methods. Silhouette width with different *p* values were calculated at each group number of the hierarchical classifications between 2 and 20, then mean silhouette widths were compared across group numbers, *p* values and classification methods.

Computations were carried out by the R software (R Core Team 2017) using the cluster package (Maechler et al. 2017). Programme codes for silhouette width using generalized mean and for generating artificial data set are available in the Supporting Information.

## Results

In all cases we inspected, except those with randomized clustering, within the same classification mean silhouette width (MSW) decreased with increasing *p*. With artificial data, when the point pattern was random, there were only up to five misclassified objects, for *p* values up to zero there were two or three misclassified objects, while for higher *p* values there were five or six ones (Fig. 1). Despite the low misclassification rate, MR decreased from 0.73 at *p* = −∞ to 0.181 at *p* = ∞. Misclassified plots were situated near the border between the two clusters. When the separation and compactness were moderate (Fig. 2), for *p* = −∞ and *p* = −2 there were two and one misclassified objects, respectively, otherwise all plots were correctly clustered with higher *p* values. There were no misclassifications at all when points were clustered into four aggregations (Fig. 3); however, MSW decreased from 0.96 to 0.77 with increasing *p*. When the same points were split into two clusters instead of their true aggregations, misclassification rate (MR) did not change but MSW decreased more steeply, reaching 0.249 with *p* = ∞ (Fig. 4). When two, well separated and compact groups were of different sizes, MR and MSW decreased as *p* increased. With *p* = −∞, there were no misclassification, and MSW was 0.92 (Fig. 5). With increasing *p* misclassified objects appeared gradually in the larger cluster near the border of the two clusters but they were not abundant until *p* = 3. However, with *p* = ∞ as high as 33% of all objects were indicated misclassified, all belonging to the larger group, and MSW were 0.202. In case of parallel groups, all objects were considered correctly classified with *p* < 0 (Fig. 6). From *p* = 0 the MR increased from 0.03 to as high as 0.6 at *p* = ∞. At *p* = −∞ MSW was 0.84, with *p* = 1 it was 0.124, while with higher *p* values MSW was near 0 indicating an unsatisfactory classification. Objects in marginal position in the point clouds tended to be identified as misclassified. With concentric groups, the inner, compact group was considered perfect regardless the *p* parameter (Fig. 7). However, the assessment of the outer group varied greatly. With *p* = −∞ all objects were deemed correctly classified. As *p* raised, the number of misclassified objects in the outer group increased, too. With *p* = 0 misclassified plots gave 23% of the total data set which means 46% of the outer group. From *p* = 1 and higher all objects in the outer group were considered misclassified, thus the data set consisted of a perfect and a totally bad cluster together giving 50% correct classification rate. Along the gradient in the parameter value, MSW decreased from 0.92 (*p* = −∞) to 0.153 (*p* = ∞). When clustering of objects was random, MR and MSW showed variable response along increasing *p* value. In case of four point aggregations but randomly permuted cluster labels MSW increased with increasing *p* parameter, while MR showed irregular response (Fig. 8). However, these silhouette width values still indicated poor clustering, since MSW ranged between −0.308 and −0.0211, while MR between 0.70 and 0.81.

**Figure 1.**
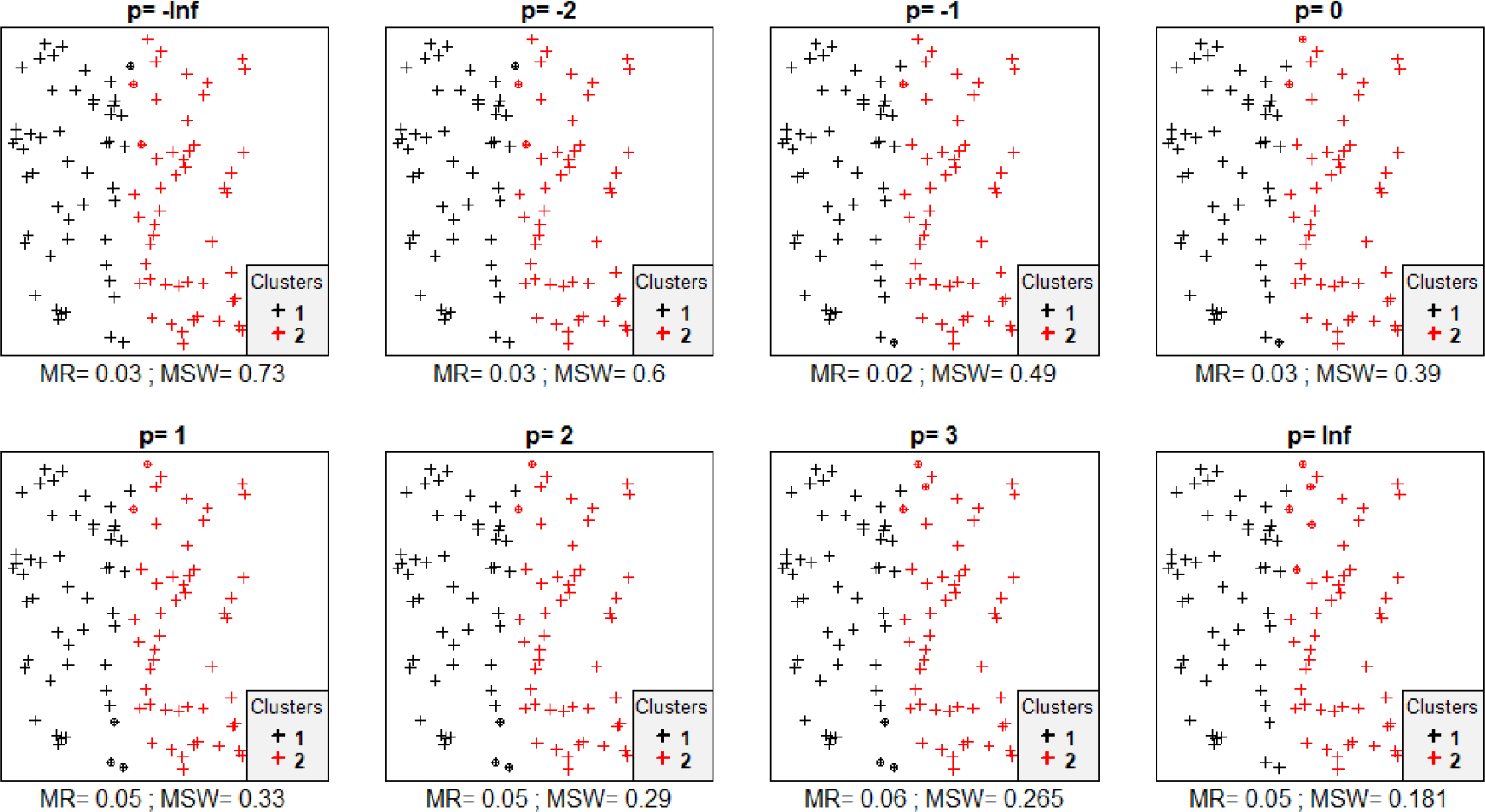
Silhouette width patterns of objects grouped into two clusters with low separation and low compactness. MR = misclassification rate; MSW = mean silhouette width; misclassified objects are circled

**Figure 2.**
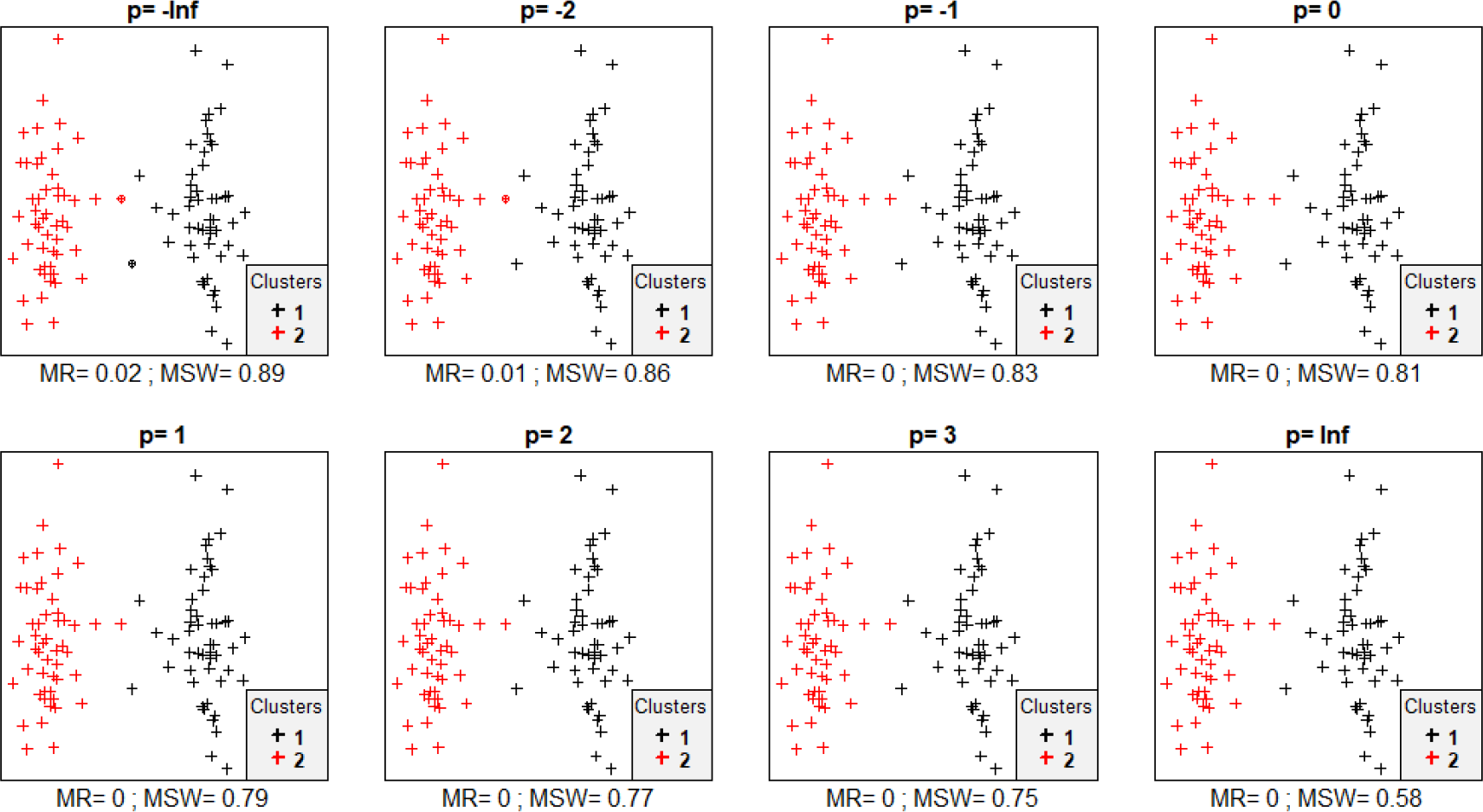
Silhouette width patterns of objects grouped into four clusters with moderate separation and moderate compactness. MR = misclassification rate; MSW = mean silhouette width; misclassified objects are circled

**Figure 3.**
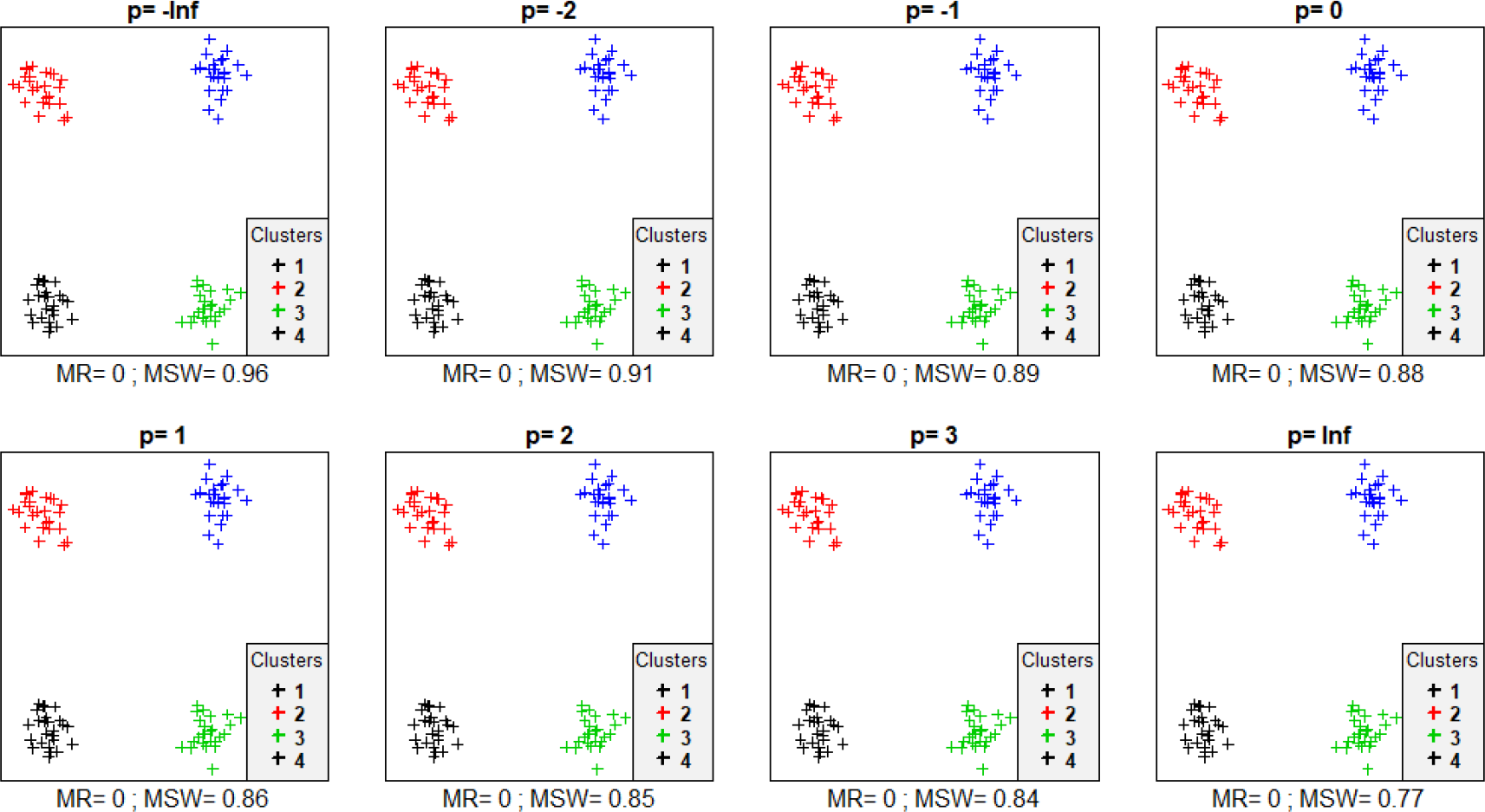
Silhouette width patterns of objects grouped into four clusters with high separation and high compactness. MR = misclassification rate; MSW = mean silhouette width; misclassified objects are circled

**Figure 4.**
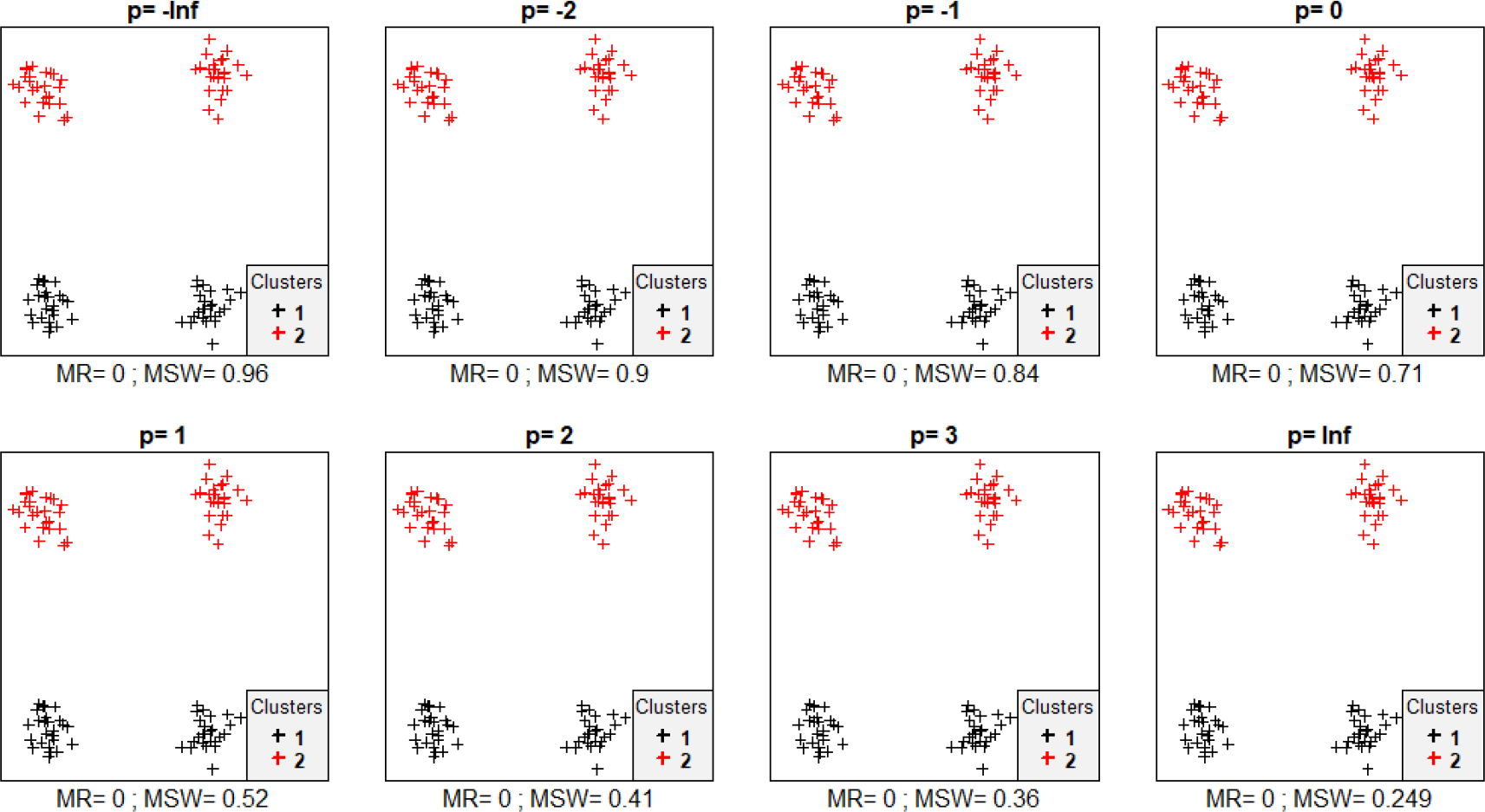
Silhouette width patterns of objects in four aggregates grouped into two clusters with high separation and low compactness. MR = misclassification rate; MSW = mean silhouette width; misclassified objects are circled

**Figure 5.**
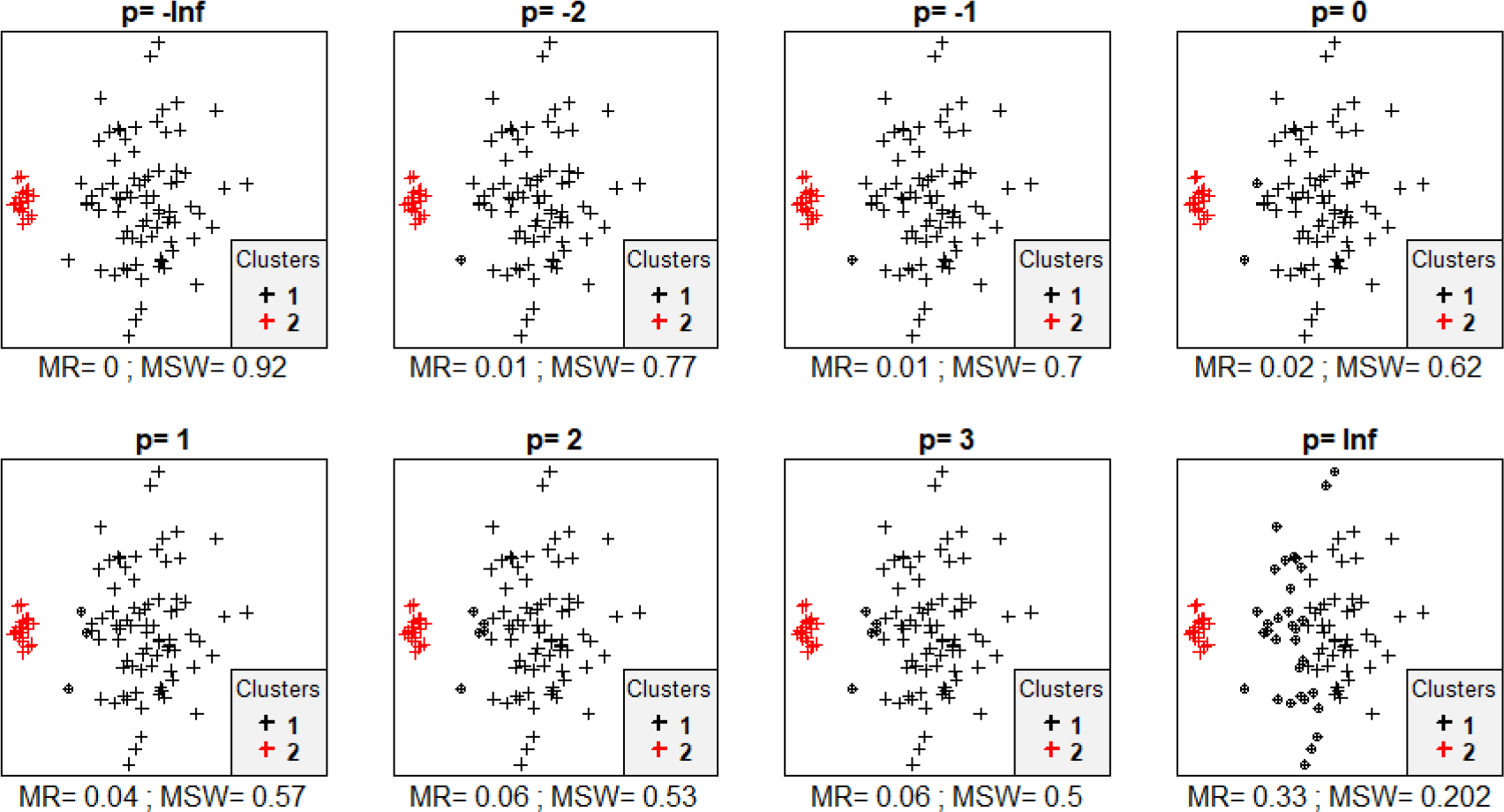
Silhouette width patterns of objects in four aggregates grouped into two clusters with high separation, high compactness and different size. MR = misclassification rate; MSW = mean silhouette width; misclassified objects are circled

**Figure 6.**
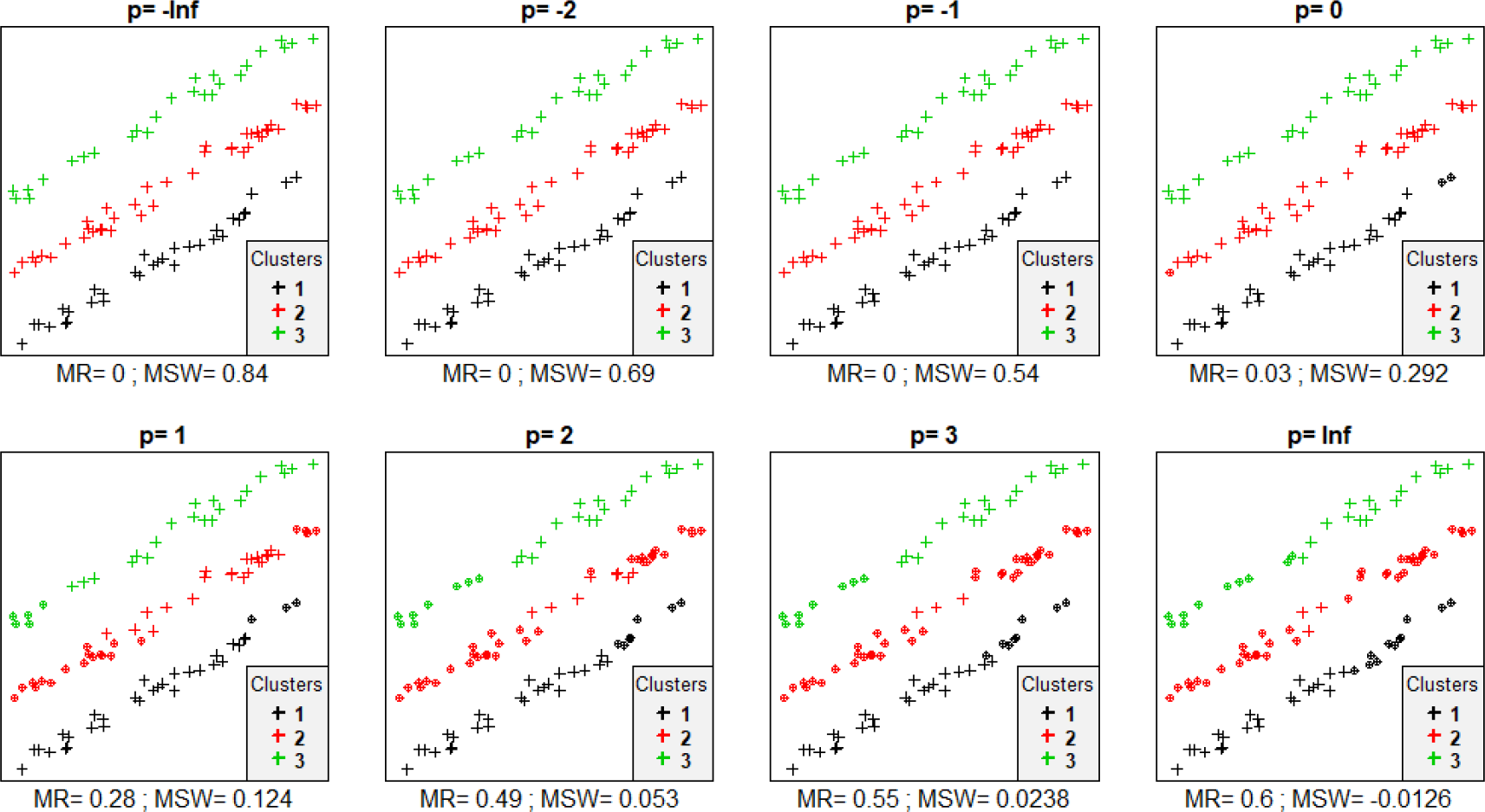
Silhouette width patterns of objects grouped into three, parallely situated clusters with high separation and low compactness. MR = misclassification rate; MSW = mean silhouette width; misclassified objects are circled

**Figure 7.**
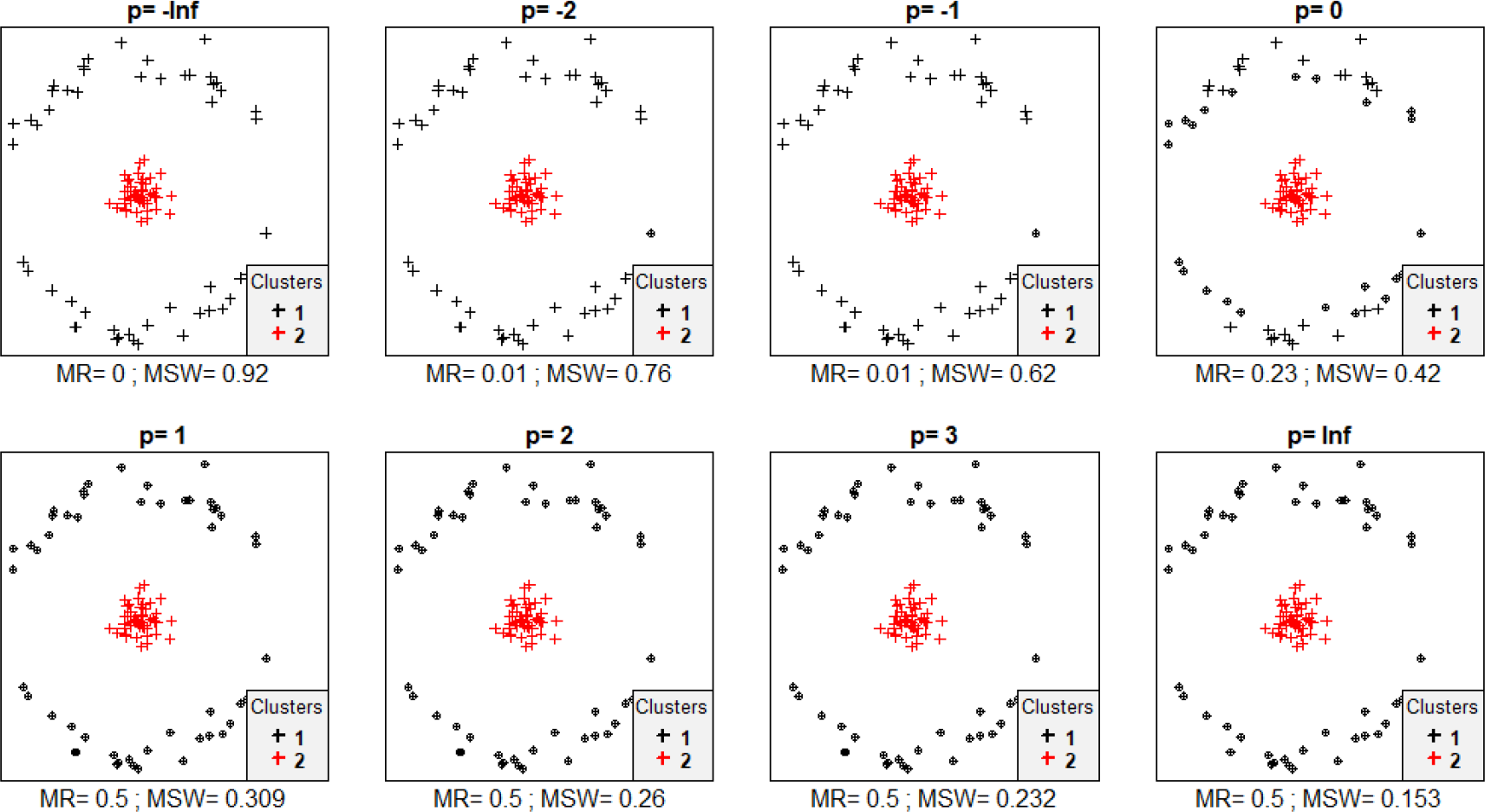
Misclassification patterns of objects grouped into two concentric clusters with good separation – an outer one with low compactness and an inner one with high compactness. MR = misclassification rate; MSW = mean silhouette width; misclassified objects are circled

**Figure 8.**
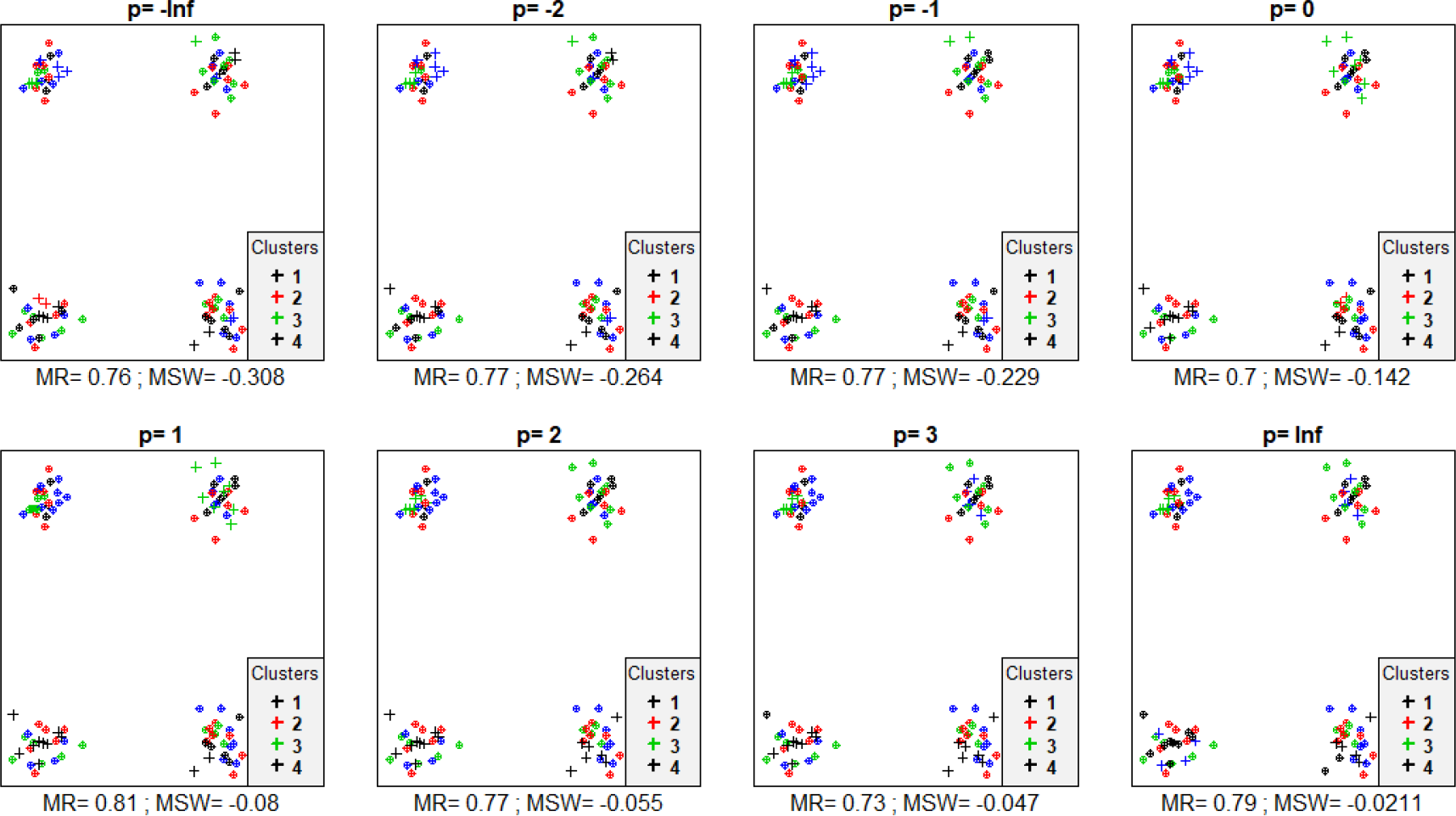
Silhouette width patterns of objects in four aggregations but with cluster assignments permuted randomly. MR = misclassification rate; MSW = mean silhouette width; misclassified objects are circled

Similarly to the simulated data, with the Iris data set, misclassification rate increased with increasing *p* parameter (Fig. 9 & 10). The minimum was 0.087 with p < 0, the maximum was 0.200 at *p* = ∞. MSW decreased from 0.71 to 0.237. *I. setosa* was perfectly separated from the other two groups, since none of its members obtained negative silhouette width with any value of *p*. At the area where *I. versicolor* and *I. virginica* overlap there were misclassified objects according to all values of *p*. However, with increasing *p*, *I. versicolor* individuals at the opposite end of the point cloud of the cluster, i.e. closer to points of *I. setosa*, also tended to seem misclassified.

**Figure 9.**
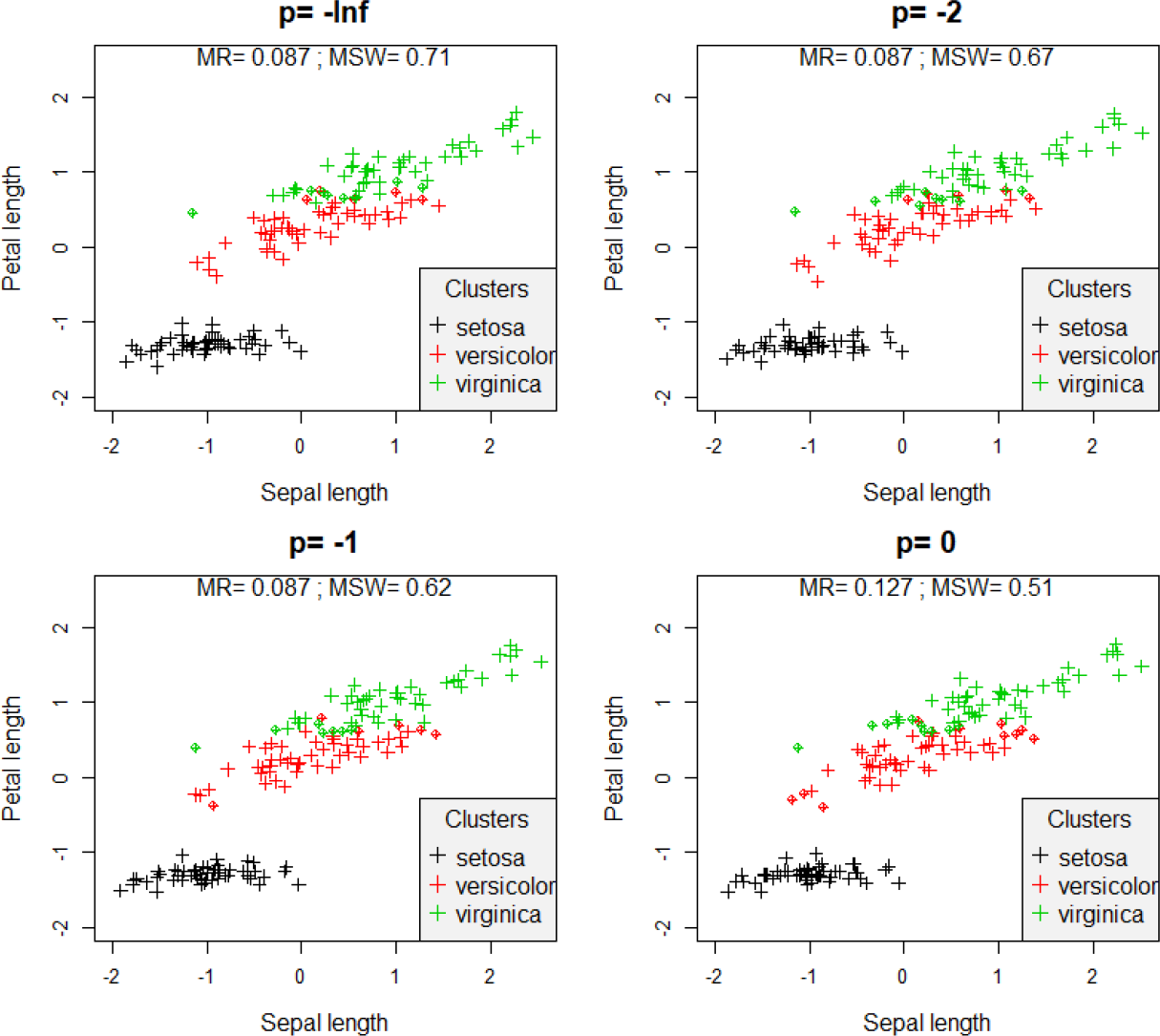
Silhouette width patterns of the Iris data set using sepal length and petal length variables after standardization to mean = 0 and standard deviation = 1 with *p* ranging from −∞ to 0. MR = misclassification rate; MSW = mean silhouette width; misclassified objects are circled

**Figure 10.**
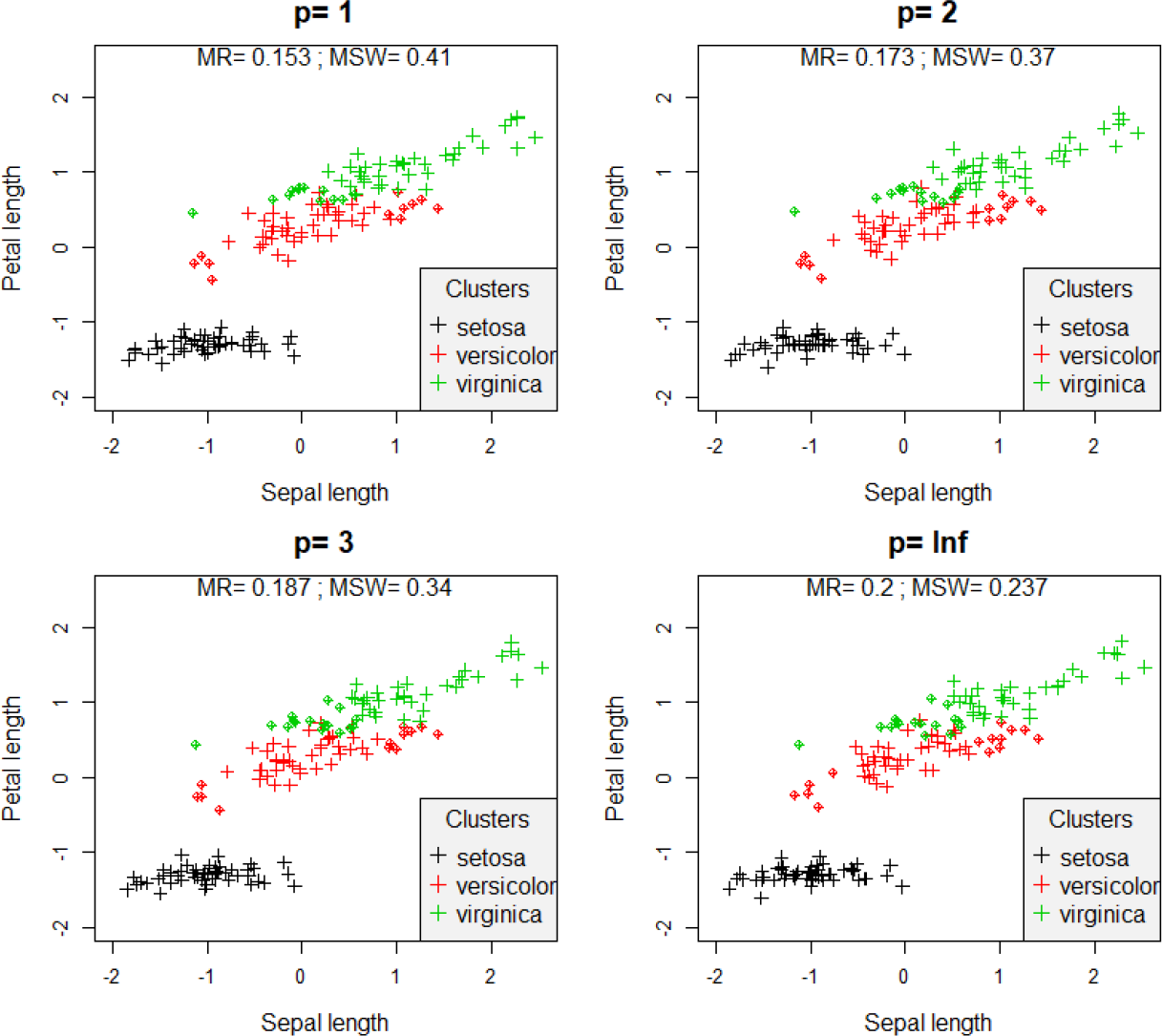
Silhouette width patterns of the Iris data set using sepal length and petal length variables after standardization to mean = 0 and standard deviation = 1 with *p* ranging from 1 to +∞. MR = misclassification rate; MSW = mean silhouette width; misclassified objects are circled

With all classification methods average silhouette width decreased with increasing the *p* parameter (Fig. 11). Using single linkage and *p* = −∞, MSW decreased monotonically with increasing number of clusters, while with higher *p* it first decreased until a minimum between 10 to 30 clusters then increased with the number of clusters. With group average and complete linkage lower (typically negative) *p* values resulted in MSW curves decreasing monotonically, while higher *p* values did not show clear trend. Nevertheless, the effect of changing the *p* value was significantly stronger on MSW when the data set was classified by the single linkage method than with the other two. When methods were compared, with *p* = −∞, single linkage obtained the highest MSW, followed by group average, and finally complete linkage – although, the latter two performed very similarly (Fig. 12). With *p* = 1, group average was slightly better than complete linkage, while single linkage obtained by far the lowest silhouette widths. With *p* = ∞, group average and complete linkage possessed similarly high average widths, while single link seemed again much less efficient at all cluster levels.

**Figure 11.**
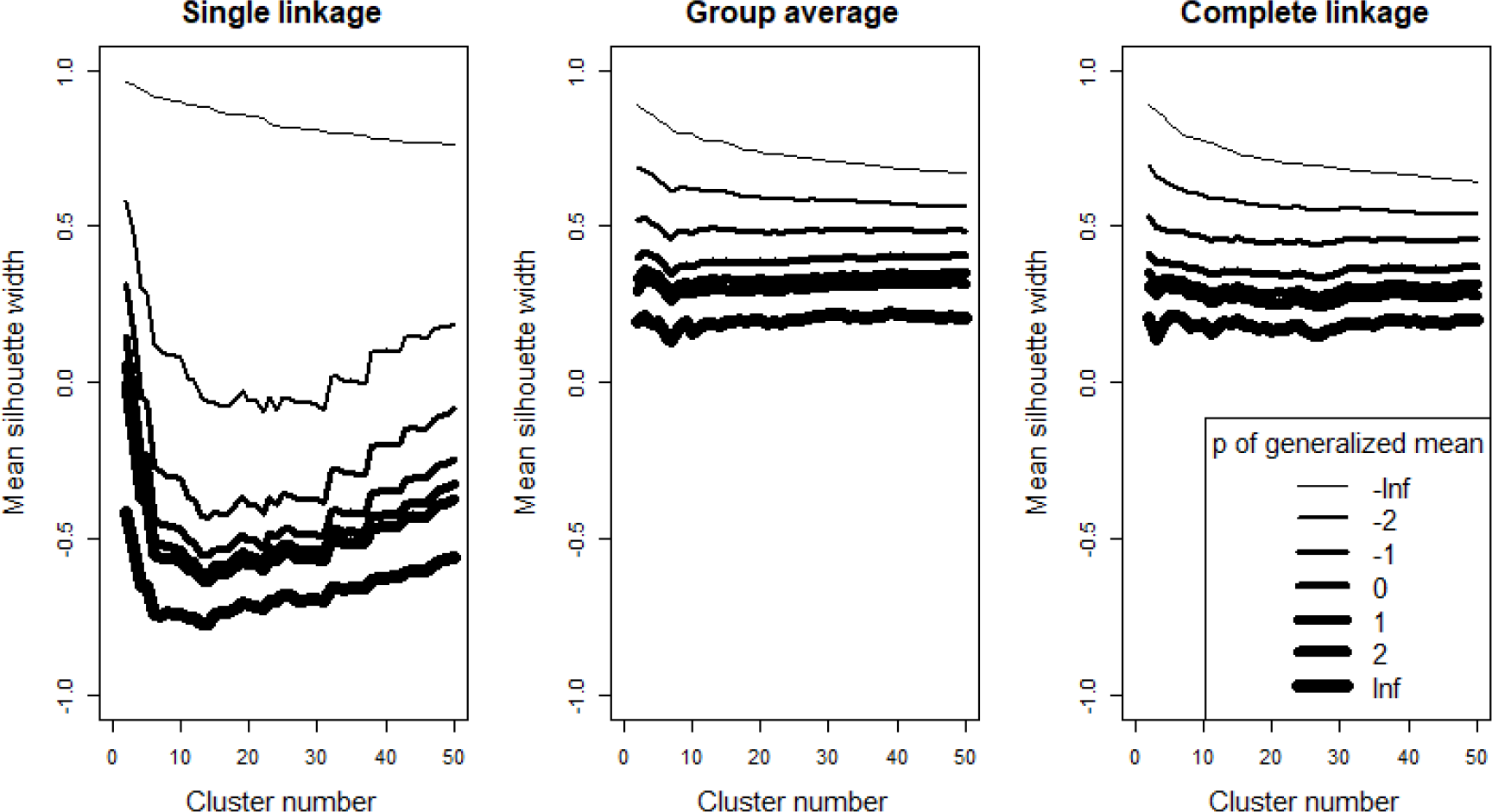
Comparison of average silhouette widths calculated with different *p* values on classifications with different methods and cluster numbers – a comparison between *p* values separating the effect of classification methods

**Figure 12.**
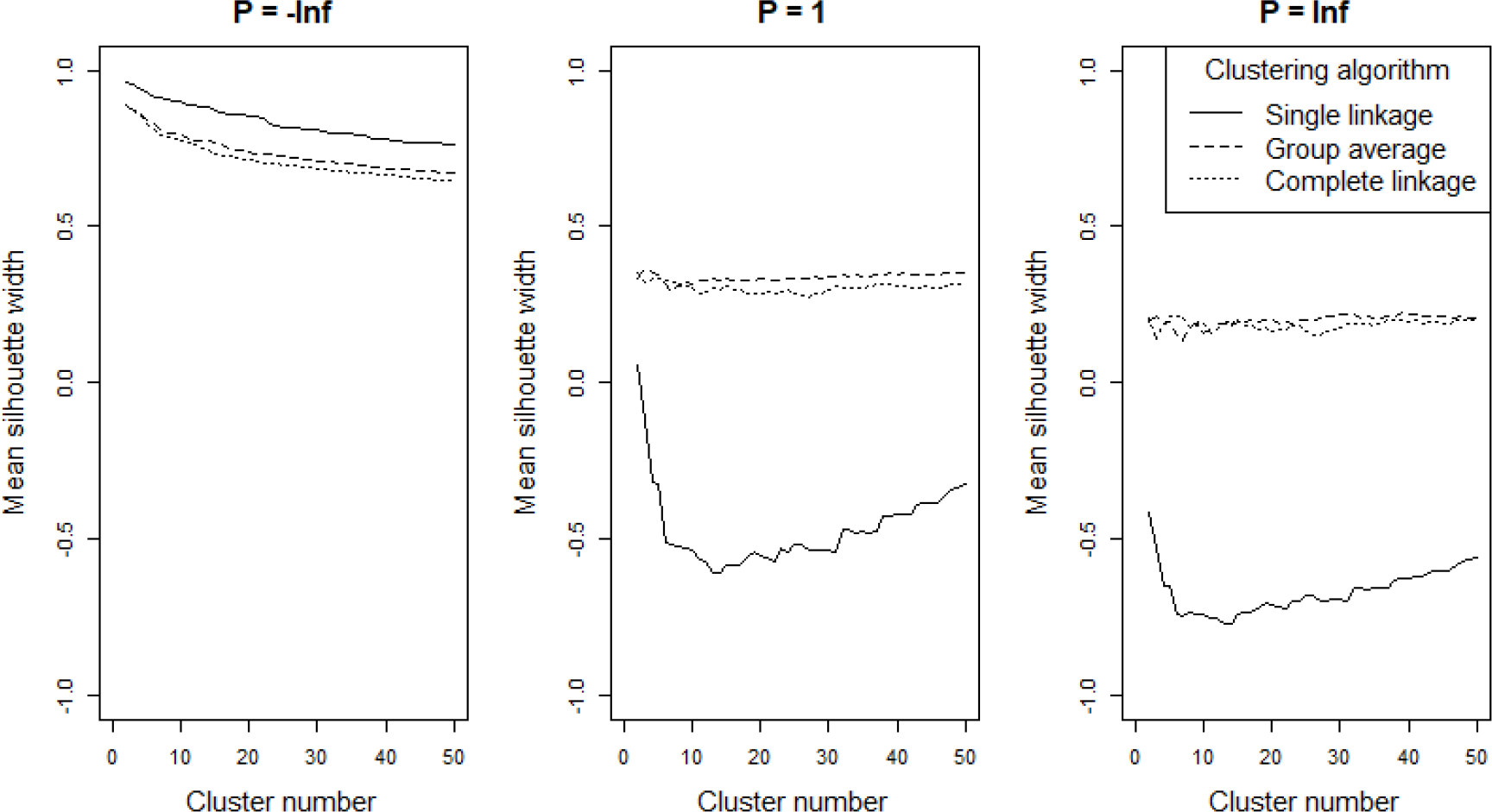
Comparison of average silhouette widths calculated with different *p* values on classifications with different methods and cluster numbers – a comparison between classification methods, separating the effect of *p* values

## Discussion and Conclusions

The results supported our expectation about the behaviour of the silhouette method using the generalized mean. Both artificial data and the Iris data set showed that cluster compactness plays less and less significant role in the assessment of classification validity with decreasing *p* parameter value. With *p* << 0 clusters are assessed mainly on the basis of connectedness and separation criterion, which in the extreme case (*p* = −∞) means the relativized difference between the minimal distances of objects belonging to the same cluster and to different clusters, while distances from other members of the same and the neighbour cluster are completely disregarded. As we increase the *p* parameter, more importance is attributed to more distant objects within and between clusters, i.e. to the compactness criterion.

When classifications were intuitively efficient from some aspect, mean silhouette width decreased, and in several cases misclassification rate increased, with increasing *p* value. In other words, these classifications tended to seem less and less efficient as the compactness criterion was attributed more and more importance. Nevertheless, with randomized assignment of objects to clusters the opposite tendency, that is, increasing mean silhouette width with increasing *p* value, was also detected in some cases (only one example shown, Fig. 8). Notably, across all tests, MSW with *p* = −∞ ranged from −0.308 to 0.960, while with *p* = ∞ this interval was much narrower, between −0.021 and 0.770. Conclusively, the relationship between mean silhouette width and the *p* parameter value is highly dependent on the data set and on the classification but with lower *p* values MSW varies on broader range. Therefore, special caution is advised if MSWs obtained with different *p* values are compared. Probably such comparisons are valid only if, instead of the raw MSW, their standardized difference from the expected value given an appropriate null model is used (Handl et al. 2005).

With different values of the *p* parameter silhouette width considers different clustering strategies effective. As it was expected, low *p* values prefer algorithms which disregard cluster compactness, e.g. single link, while with high *p*, procedures resulting in spherical clusters (e.g. group average, complete linkage) are deemed better. In the comparison of classification methods in the view of the generalized silhouette width, group average and complete linkage behaved similarly efficiently across different *p* values and cluster numbers.

There are many other cluster validation indices that combine cluster separation and compactness (Handl et al. 2005; Vendramin et al. 2010), however silhouette width is the only one that evaluates individual objects. Generalized mean instead of arithmetic mean (or minimum or maximum) could be used in other indices combining the separation and the compactness criteria. Similar examples are already shown by Bezdek & Pal (1998) for the generalization of the Dunn index.

Ideally, ‘good’ clusters should show a spherical shape given that the variables and their weight in the analysis are selected appropriately. However, in reality the selection of variables is constrained by serious practical limitations, and usually there is no objective recommendation on the method for weighting. Therefore, natural objects frequently show non-spherical shapes in the multidimensional space of the analysis. In such cases, a cluster validity measure with a preference towards spherical shape can evaluate cluster quality too rigorously. When it is not reasonable to expect spherical clusters but only their connectedness and separation is relevant, setting *p* to negative values to assess the fit of objects into the classification can be a solution. We especially advise to calculate silhouette width with different values of *p*. In this way, a new dimension of methodological decisions referring to cluster compactness can be involved into the assessment of classifications (Lengyel et al. 2018). However, we recall that raw silhouette widths with different parameterization may not be directly comparable, since with lower *p* values widths vary on broader range. Hence curves of average silhouette width with different *p* values along number of clusters should be viewed as different indices which are ordered by sensitivity to compactness, and no ‘optimal *p* value’ should be sought for empirically.

## Acknowledgements

A.L. was supported by the National Science Centre, Poland, through the POLONEZ fellowship programme (grant 2016/23/P/NZ8/04260). The Authors state that they have no conflict of interest.

## Authors’ contributions

A.L. developed the idea and the methodology, wrote the scripts, conducted data analysis and wrote the manuscript, Z.B.D. developed the idea, reviewed literature, commented on the results, and improved the manuscript.

## Data accessibility

Scripts for calculating generalized mean and generating specific point patterns are enclosed in the Supporting Information. Iris data set is available from the vegan package of R.

